# Aggregation tendencies of a female *Vaejovis carolinianus* population on France mountain, Tennessee

**DOI:** 10.1101/058362

**Authors:** Bob A. Baggett, Hannah E. Ritter, Margaret E. Melendez

## Abstract

France Mountain is located within Overton County which is part of the Upper Cumberland region of Middle Tennessee contains only one species of scorpion, also the only species native to the state, *Vaejovis carolinianus*. This species is poorly researched, and little is known about its life history and ecology. The objectives of this study were to determine if female *V. carolinianus* prefer to form aggregations under large cover objects or seek out retreat sites where they would be the sole occupant. Based on past research, we hypothesized that females would form aggregations under large cover objects instead of being the sole occupant. A total of 32 scorpions were captured during August and September 2014. During laboratory trials, three scorpions were placed in a plywood tray containing sand substrate and three equally-sized large ceramic tiles. The scorpions were left undisturbed for a 24-hour period, after which the tiles were lifted to check for aggregations. Each group of three scorpions was in the research tray for three consecutive days. Four rounds of aggregation trials were conducted, and t-tests as well as linear regression were used to analyze the data collected. The results of the t-test showed that female *V. carolinianus*, when given a choice of similarly-sized large cover objects, would select a retreat site where they were the sole occupant; therefore, our initial hypothesis was rejected. Linear regression found relative humidity affected aggregation occurrence, for the largest number of aggregations occurred when the relative humidity was between 67 and 73%.

## INTRODUCTION

How and why organisms select particular habitats have long been important topics in ecology (Huey 1991), for habitat selection by mobile organisms can have significant consequences for an animal’s biology, from defining extent of environmental interactions to conferring significant fitness advantages (Goldsbrough et al. 2004). Factors contributing to habitat selection include: (1) environmental factors (temperature, precipitation, soil type, etc.); (2) availability of shelters, food, nest sites, and mates; (3) degree of intra-and interspecific competition; and (4) risks of predation, parasitism, and disease (Hodara and Busch 2010).

Retreat sites are locations selected by organisms for safety from predators, accessibility of food, and/or appropriate thermal or humidity regime (Langkilde and Shine 2004). Many animals spend long periods of time within the shelter of their retreat sites, and individuals are often philopatric to such sites (Croak et al. 2008). Choice among retreat sites may be based on scent cues from the same or other species, conditions offered by the site (i.e., moisture content, protection from sunlight), or physical structure of the retreat site itself. Availability of suitable retreat sites may have strong impacts on individual fitness as well as population viability. Accordingly, animals select retreat sites nonrandomly and often use multiple criteria to prioritize suitable sites (Croak et al. 2008).

In many species, individuals aggregate in response to a variety of cues (e.g., thermal, hydric, nutritional, reproductive, predator-based) and obtain an array of potential benefits from such behavior (Aubret and Shine 2009). Aggregation is defined as two or more organisms sharing a cover object (assumed to be aware of each other’s presence), and the organisms may or may not be in physical contact with one another (Gregory 2004).

Of the 98,000+ arachnid species currently described, less than 0.06% are social (Rayor and Taylor 2006). Scorpions, for the most part, are solitary and sedentary organisms that prefer microhabitats colonized by their prey. Intraspecific and heterospecific coexistence has been observed in several scorpion species, producing different levels of sociability and aggregation (Lira et al 2013). “Social phases” are normally limited to mating and maternal care of offspring from birth to shortly after the first instar (Mahsberg 1990). In some species, unrelated individuals occur together in the field, for mature males and females may share a burrow or the same ground cover during the mating season (Polis and Lourenço 1986). Other species tend to aggregate in larger groups; for example, *Centruroides sculpturatus* can be found in groups of up to 30 individuals during the winter (Polis and Lourenço 1986). Kaltsas et al. (2009) distinguishes two evolutionary routes regarding aggregation in scorpions: (1) the familial route (aggregation of relatives) and (2) the communal/parasocial route (aggregation of nonrelatives). Regardless of evolutionary route, it is believed that scorpions aggregate in one place merely because conditions are favorable (Abushama 1964).

Members of seven scorpion families reside in North America; however, only one family, Vaejovidae, is endemic to North and Central America. Vaejovidae is comprised of 23 genera and 200 species (Rein 2015). Vaejovids occur in most terrestrial habitats. Some burrow, while others seek shelter under rocks, fallen logs, or leaf litter. Two species of scorpions are found in Tennessee, but only one, *Vaejovis carolinianus*, is native to the state. *Vaejovis carolinianus* has many common names, but two most commonly used are Plain Eastern Stripeless Scorpion and Southern Devil Scorpion. Reddish to dusty-brown in color, *V. carolinianus* may reach up to 6.5 cm in total length with a lifespan of seven to eight years (Benton 1973). *Vaejovis carolinianus* can be found in the southeastern United States. It commonly resides along edges of moist forest habitats, which might provide shading and close proximity to food and water resources (Croak et al. 2012); it is almost never found in prairie grasslands, large open fields, mountain meadows, or recently altered areas such as strip mine spoil banks (Benton 1973). During the day, they are usually located under rocks, leaf litter, or the bark of dead trees. This scorpion also occurs around human dwellings if sufficient cover is available. Unlike other species in its family, *V. carolinianus* is not a strong burrower, and it uses retreat sites with natural spaces or crevices (Benton 1973). *Vaejovis carolinianus* is most active above 25°C and is sluggish in colder temperatures (Elston 2006). *Vaejovis carolinianus* is viviparous with a gestation period of one year. Parturition (Figure 4) usually occurs in mid-to late August (Taylor 1970).

Male and female *V. carolinianus* have different habitat preferences (Benton 1973; Dobbe 1999). Adult females typically occur under and among rocks, as well as at the base of dead standing trees. Males and immature scorpions occur most commonly in leaf litter, as well as under the bark of dead trees (preferably pine). Mating occurs in late August; otherwise, adults do not usually interact (Polis and Sissom 1990; Dobbe 1999). However, up to nine females have been found hiding under the same rock (Benton 1973). Such aggregations are not considered social groups, but instead likely represent groups of individuals seeking similar resources. The proximate basis of aggregation formation may involve ecological and/or behavioral factors. Ecological factors include aggregations forming around limiting resources (water, food, or preferred habitat) that have a clumped or patchy spatial distribution, while behavioral factors involve individuals seeking conspecifics, sometimes referred to as conspecific attraction (Hodge and Storfer-Isser 1997).

The objective of this study was to determine if, given an abundant selection of large cover objects, female *Vaejovis carolinianus* would aggregate. Our hypothesis was that the females would form aggregations of at least two individuals more than 50% of the time.

## MATERIALS AND METHODS

### Study Sites and Scorpion Collection/Maintenance

A total of 35 *Vaejovis carolinianus* (26 females, 6 males, and 3 juveniles) was captured from four sites near Pleasant Shade Lane on France Mountain in Tennessee’s Upper Cumberland region during August through October 2015. The first site was a campfire pit located along an electric line access (36.241653° N, 85.236558° W, altitude 399.90 m). The entrance to the electric line access was 4.99 km from the intersection of Pleasant Shade Lane and Dry Hollow Road. Site two was a large rock (we called it Hannah’s Rock) located at the intersection of the electric line access and an unnamed Jeep trail (36.242731° N, 85.235875° W, altitude 405.69 m). Site three was a small area (36.238292° N, 85.239536° W, altitude 398.37 m) on a logging road, the entrance to which was located 4.67 km from the intersection of Pleasant Shade Lane and Dry Hollow Road. The logging road site had been heavily impacted by both timber cutting and rock collection. Finally, site four was an approximately 50 m stretch of roadside (36.237569° N, 85.239464° W, altitude 401.12 m) along Pleasant Shade Lane just before access to the logging road.

Most scorpions were captured during the day by turning over rocks and logs along the edges of moist forests and in road cuts. The remaining scorpions were captured at night by walking along Pleasant Shade Lane and illuminating the edge of the road with a UV-A light. Upon capture, scorpions were placed together in a large plastic container for transport back to Tennessee Technological University (TTU). At TTU, scorpions were individually placed in a 7.5-cm x 6.3-cm Ziploc^^®^^ plastic bag to be weighed (in mg) and measured (total body length minus the telson and aculeus, in cm). Sex of each scorpion was determined by examining the pectines, which are comb-like sensory organs located ventrally; these are larger on males and contain more teeth. Total length, weight, sex, and a unique identifier number were recorded in an Excel spreadsheet. Scorpions were tagged dorsally with a small queen bee marker tag and placed individually in a 17-cm x 11-cm x 5.8-cm plastic container containing a 1-cm deep layer of sand substrate with a square (~4 cm^2^) of moistened paper towel for water and cover. Scorpions were fed one banded cricket, *Gryllodes sigallatus* of the same size as the scorpion or smaller, every 7-10 days while in captivity. Paper towels were daily to maintain moisture and examine for mold. Paper towels showing any evidence of mold were immediately replaced.

### Laboratory Procedures

Experimental trials started 17 September 2015. All trials were conducted in Room 409A of Pennebaker Hall at Tennessee Technological University. This room was selected due to its limited access, plus its ambient temperature was similar to outside temperature. Ambient temperature ranged between 17.6° and 23.9° C. All trials were conducted in plywood trays measuring 0.914m x 0.914m (0.836m^2^) and 15.24 cm high. The interior of the tray was lined with wax paper to prevent the scorpions from crawling up the side and escaping. Trays were filled with 4 cm of playground sand, and three ceramic tiles (929 cm^2^ each) were placed in a triangular pattern in each tray. A light using a 40 watt incandescent bulb was placed directly over each tray, and the light was controlled by a timer to establish a 14 hour light; 10 hour dark daily cycle. This cycle was selected based on average sunrise/sunset patterns in late summer and early fall in Tennessee. Scorpions were kept in their individual containers in this same room, so no acclimation time was required prior to testing.

Aggregation was defined as two or more scorpions sharing a cover object regardless of whether they were in actual physical contact with one another (Gregory 2004). Three female scorpions were placed in the center of the tray at the beginning of the light cycle and allowed to remain undisturbed for one 24-hour light:dark cycle. At the end of the dark cycle, the tiles were lifted and each scorpion’s location was recorded. An aggregation would be recorded if scorpions were within 5 cm of each other, and it was also noted whether the aggregation involved two or three scorpions. Additional data collected included time, ambient temperature, humidity, and temperature under each tile.

Once data was recorded, scorpions were pulled from the trays and tiles were rotated in a clockwise manner. The scorpions were then returned to the center of the tray and left undisturbed for another 24-hour cycle. Each group of three scorpions remained in the trays for three consecutive days. At the end of the three-day trial period, scorpions were removed, tiles were wiped clean, and another group of three scorpions was placed in the tray. Each scorpion went through four rounds of aggregation trials, and no two scorpions were grouped together for more than one round.

### Data Analyses

Aggregation tendency was analyzed using a *t*-test comparing each round of data with an expected aggregation value. No aggregation was assigned a value of 0, an aggregation of two scorpions were assigned a value of 2, and an aggregation of three scorpions was assigned a value of 3. Daily aggregation values were added together for a round score, and this score was compared to an expected round score of 6 (assuming that an aggregation of at least two scorpions would occur each day). T-tests were conducted to compare actual aggregation scores to expected aggregations scores for the complete trial period as well as for each round. An additional t-test was used to determine if there was a significant difference between the temperature under the tile selected for aggregation and the other tiles. Linear regressions were used to determine if ambient temperature or relative humidity had any effect on aggregation.

## RESULTS

Figure 1 shows the breakdown of the total number of aggregations for the entire trial period. T-tests were used to compare actual aggregation scores to expected aggregation scores for each round as well as for the total trial period. For Round 1, significance was found between the two scores (t = −20.725, df = 27, *P* < .001). The t-test for Round 2 also found significance between actual aggregation scores and expected aggregation scores (t = −6.424, df = 16, *P* < .001). T-tests for Rounds 3 and 4 also found significant differences between actual aggregation scores and expected aggregation scores (t = −4.925, df = 8, *P* = .001), (t = −14.000, df = 4, *P* < .001), respectively. The t-test for the entire trial period found a significant difference between actual aggregation scores and expected aggregation scores (t = −18.595, df = 58, *P* < .001). Finally, no significance was found between the temperature under the tile selected for aggregation and the temperature under the other tiles (t = .030, df = 15, *P* = .976).

**Figure 1.**
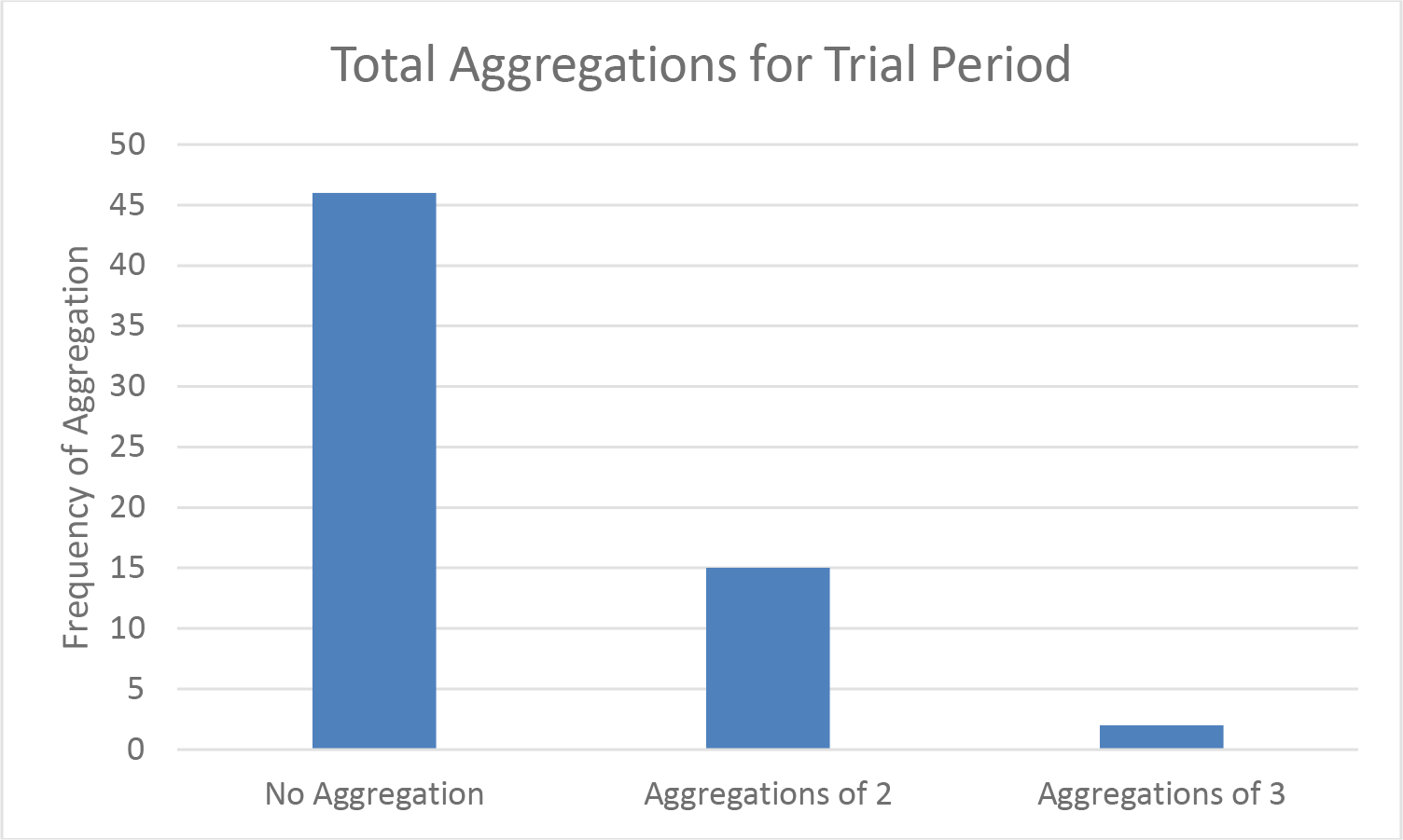
Column graph showing the total number of aggregations over the entire trial period.

A linear regression used to determine if aggregation was affected by ambient temperature found no significance (r^2^ = .047, F_1,29_ = 1.417, *P* = .244). However, linear regression used to determine if aggregation was affected by relative humidity found significance (r^2^ = .289, F_1,29_ = 11.781, *P* = .002). A scatterplot (Fig. 2) was created to show the number of aggregations occurring at the recorded relative humidity levels.

**Figure 2.**
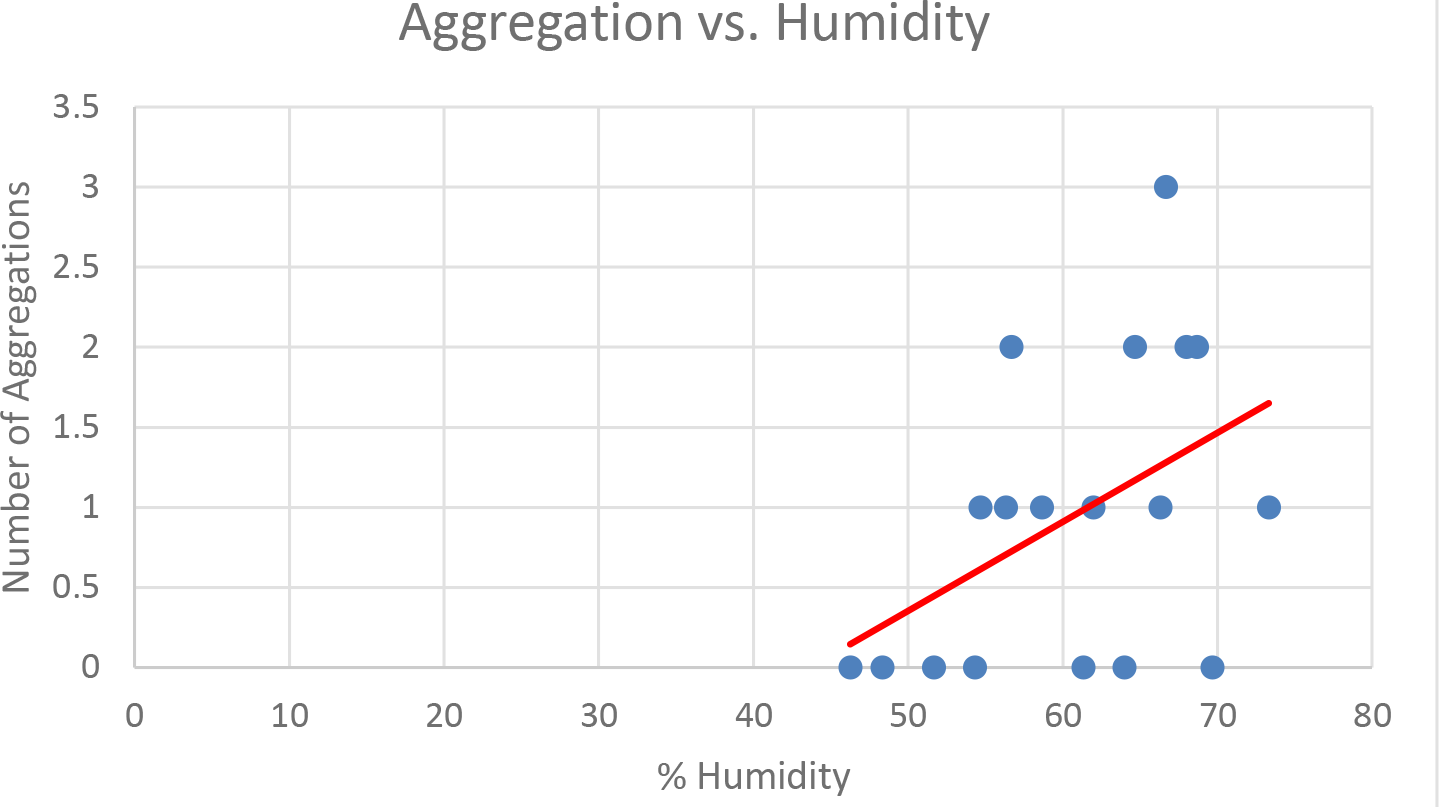
Scatterplot showing the number of aggregations occurring at various humidity levels.

## DISCUSSION

Animal aggregations differ from social groups since aggregations represent groups of individuals engaged in individual pursuits coincidentally in the same vicinity (Hodge and Storfer-Isser 1997). Lancaster et al. (2006) concluded that banded gecko (*Coleonyx variegatus*) aggregations occur in order to reduce evaporative water loss by increasing retreat site humidity. Two species of web-building spiders (*Hypochilus thorelli* and *Achaearanea tepidariorum*), aggregate for behavioral reasons. These species will attach their webs to their neighbors’ webs (Hodge and Storfer-Isser 1997). Similarly, scorpions likely aggregate in one place merely because conditions there are preferable (Abushama 1964). A study conducted in Kentucky found as many as nine *V. carolinianus* females aggregating under one large rock in January, and it was common for two or three to be found together (Benton 1973). It was assumed that males did not aggregate (Benton 1973), but males are seldom found. These aggregations may occur because of a lack of suitable habitat for overwintering, so scorpions tended to gather under cover objects that were more protective, e.g., deeper into the soil or underneath heavier leaf litter (Benton 1973). Aggregations occur in other scorpion species as well; for example, McAlister (1966) determined favorable microhabitats led to aggregations in a Texas population of *Centruroides vittatus*. An aggregation of nine *C. vittatus* scorpions was in the dry interior of an oak stump during a heavy rain, and aggregations were found in moist sand beneath logs during intervals of cold weather (McAlister 1966).

Based on the t-test results, we had to reject our initial hypothesis. Our aggregation trials indicated that female *Vaejovis carolinianus* scorpions may aggregate, but prefer to select retreat sites not occupied by other scorpions. All three tiles used in each tray were large tiles of equal surface area since a previous study (Baggett 2015) found that scorpions select large cover objects as retreat sites. During the Baggett (2015) study, tiles of three different surface areas were used, and this somewhat biased the aggregation trial since the scorpions involved preferred large cover objects. The current study eliminated such bias. Baggett’s 2015 study also controlled for humidity and ambient temperature by conducting all trials in an environmental controller, whereas the current study allowed for daily variations in ambient temperature and relative humidity. Based on our linear regression results, ambient temperature had no effect on aggregation preferences, but we found that aggregations were more likely to occur when the relative humidity ranged between 67 and 73%. Scorpions receive necessary water by consuming their prey, moisture in the substrate, and relative humidity (Benton 1973). Similar to the gecko aggregations found by Lancaster et al. (2006), scorpions may form aggregations in order to maintain or increase retreat site humidity. Meek (2008) found that humidity levels within retreat sites was considerably higher than external relative humidity. So, if *V. carolinianus* aggregations occur as a means of maintaining or increasing retreat site humidity, it would seem that more aggregations should occur when the relative humidity is low rather than high. Our results, however, showed the opposite.

While capturing scorpions for our aggregation trials, we found an aggregation of 3 adult female and approximately 20 juvenile scorpions. The juveniles appeared to have recently become independent of their mother’s care, and they were found under leaf litter. The females were on top of the leaf litter which was located under the Hannah’s rock site. No aggregations were found at any other site. The logging trail site was heavily impacted
by timber cutting and commercial rock collection, so the number of large rocks available was considerably lower in comparison to the other three collection sites. Future studies at this site may find an increase in aggregations under remaining large rocks, or scorpions may abandon this site in pursuit of other areas with more favorable conditions.

Time in captivity did not seem to affect the results of the aggregation trials, for all four rounds of trials showed similar aggregation preferences.

Suggestions for future studies involving *V. carolinianus* are numerous, for little is known about this seldom-researched species. In particular, more research needs to be conducted with retreat site selection; for example, studies could be conducted regarding the thermal properties of retreat sites. The actual moisture content of retreat sites should be measured, and using this data, the soil moisture preference trials could be repeated with the actual value found at the retreat sites used as the expected value during lab trials.

Other suggested studies include additional research on male *V. carolinianus,* as well as juveniles. Currently, all that seems to be known about males is that they do not aggregate, and that they prefer a different type of microhabitat as a retreat site than do females. Apparently, males will hide under the loose bark of fallen or standing dead trees, but no scorpions were found under bark during this research. Only one study was found that examined juvenile *V. carolinianus,* and this was a brief paper involving the rearing of juveniles in a laboratory environment (Taylor 1971). Future studies could examine the costs to the mother of parturition and rearing of young, as well as food preferences and growth rate of juveniles.

## ACKNOWLEDGEMENTS

Hannah, Maggie, and I thank Dr. Christopher Brown for his guidance and support throughout our research project. We also thank Sean Beddoe who assisted us in scorpion care and data collection. Additionally, we thank our families and friends who, even though they thought we were a little weird for working with scorpions, continuously offered their support and encouragement. Funding and research space for this project was provided by Tennessee Technological University.

